# Independent Online Visuomotor Control to Cursor and Target Motion

**DOI:** 10.64898/2026.06.28.735032

**Authors:** Sae Franklin, Michael Dimitriou, David W. Franklin

## Abstract

Skilled control of visually-guided reaching is fundamental for many daily activities. Visual information about hand and target position are used for movement planning and online corrections through rapid visuomotor feedback responses. Such feedback control is generally believed to implicate a single error signal, representing a difference vector between hand and target position. Here, we directly assess whether shared or independent systems serve visually-guided feedback control. We tested whether feedback gains can be independently modulated by hand/cursor and target motion through manipulating the task-relevance of each signal during goal-directed reaching. Our results demonstrate that the gains of visuomotor feedback responses to perturbed hand and target motion can be set independently of one another, at the same time, as a function of task-relevance. By dissociating feedback control of cursor and target signals, our findings support the existence of two independent visuomotor feedback pathways, revealing a more flexible neural architecture for goal-directed action.

## Introduction

Humans constantly interact with the world. We reach, grasp, manipulate and kick objects in our complex, changing environment, while avoiding distractors and collisions. In a world in which our environment can change, this means that we need to be able to re-plan and rapidly adjust our movements. Although there is evidence for planning ^1–5^ and feedforward control ^6^, we constantly rely on feedback control for accurate movements ^7^, especially given the presence of motor noise ^8,9^ and unpredictably ^10^. This often requires fine tuning of our sensorimotor control system as the task, our body, or our environment changes ^11^. During goal-directed movements, we are guided through visual feedback of the hand and the target, both to determine the appropriate movement to reach a location in the world and to produce corrective responses, such as for a moving target or to correct errors. Although voluntary corrections to visual stimuli are slow, there are rapid visuomotor responses to visual signals that occur following the presentation of a visual stimulus ^12,13^, shifts in the target location ^14^, motion of the visual background ^15^, or shifts in the visual hand location ^16–18^. These visually induced corrective responses occur relatively quickly (150 ms) after the visual stimulus and do not require conscious perception of the stimulation ^14,19^. Evidence suggest that these early components of the visually induced motor responses are involuntary ^15,20–22^. As all these motor responses to visual stimuli have similar loop delays, and there is evidence of a rapid pathway through the superior colliculus ^12,13^, it is thought that these visuomotor feedback responses might occur through a rapid feedback pathway which avoids the long processing within visual cortex.

Despite the rapid feedback nature of these visuomotor feedback responses, they are modulated according to the movement dynamics ^23–25^, the task ^26,27^, and the visual environment ^22,28,29^. The task-dependent modulation of these feedback responses is similar to those occurring in the stretch-reflexes ^30–36^ and are thought to tune the sensorimotor control system to optimise our movements in the presence of noise ^37,38^. The visuomotor responses can be tuned to the statistical properties of the environment ^22^, with an up-regulation of the feedback gains when sensory discrepancies were task-relevant, and a decrease in feedback gains when such visual motion was task-irrelevant. Moreover, these visuomotor feedback gains can be independently tuned to different states of the limb and different perturbation directions ^22,28^.

Despite the fact that rapid motor responses respond to rapid shifts in both the visual target and hand location, there is evidence that these two feedback pathways are partially independent ^39,40^. Although the difference vector model ^41^ had proposed that the visual location of the hand and target are compared to provide a single error measure guiding movement planning and control, suggesting a single feedback pathway, there are several lines of evidence against this. First, there is evidence for a visuomotor binding mechanism that confers a privileged status on visual information representing the kinematics of a moving limb, which attention must supply in the case of a visual target ^39^. Second, this model was directly tested by measuring the responses to visual perturbations over a large range of possible variations in both target and hand displacements ^40^. This work found that the rapid motor responses were consistent with two independent feedback systems. If, as our previous work suggested, these are two independent feedback control systems, then we should also be able to independently control the visuomotor feedback gains to cursor or target motion. Here we test this possibility directly by using task-relevant and task-irrelevant sensory discrepancies during reaching movements to tune the sensorimotor system and then independently probe the visuomotor feedback gains. We examine whether we can simultaneously upregulate the visuomotor feedback gains to cursor motion while downregulating the visuomotor feedback gains to target motion, and vice versa. This work directly tests if we can modulate our visuomotor feedback gain independently to hand and target motion according to task relevance.

## Results

We tested ten young adults to determine whether the visuomotor feedback response could be independently modulated to perturbations of the cursor and the target. Every participant performed 2800 forward reaching trials in each of two different visual environments. In one visual environment, during the reach, the cursor undergoes task-irrelevant lateral motion which should be ignored, while the target undergoes task-relevant lateral motion which needed to be corrected. In the other environment, these were switched so that the lateral cursor motion needed to be corrected for, whereas the lateral target motion should be ignored. We then examined whether the visuomotor feedback responses could be simultaneously tuned differentially to the motion of the cursor and the target.

### Adaptation to the visual environments

In the task-irrelevant cursor and task-relevant target condition, participants were able to improve their performance over the experiment (Fig 1). While the success rate was initially low (34.1% over the first 100 trials), over the course of the 2800 trials in this environment, participants gradually increased (t_9_=13.56; p<0.0001) their success rate, reaching 74.3 % over the last 100 trials on the second session (Fig 1A). As the success rate increased, the participants also decreased the mean duration of their movements (Fig 1B) with a significant difference from the first 100 to final 100 trials (t_9_=9.07; p<0.0001). Despite the reduction in movement duration, the peak forward speed of the movements decreased slightly (t_9_=3.02; p=0.0144) over the experiment (Fig 1C). The clear differences in the amount of lateral hand motion correcting for the visual discrepancies can be seen for trials with cursor or target motion (Fig 1D). In particular, the hand trajectories when the cursor underwent a task-irrelevant motion (smooth motion with no offset between the hand and the cursor at the end of the movement) were generally straight to the target, ending within the target location (Fig 1E). In contrast, during the intermixed trials in which the target underwent task-relevant motion (smooth motion with a final offset that needed to be corrected to complete the task successfully), the hand trajectories for each of the target motion were smoothly corrected to the final location of the target (Fig 1F).

**Figure 1.**
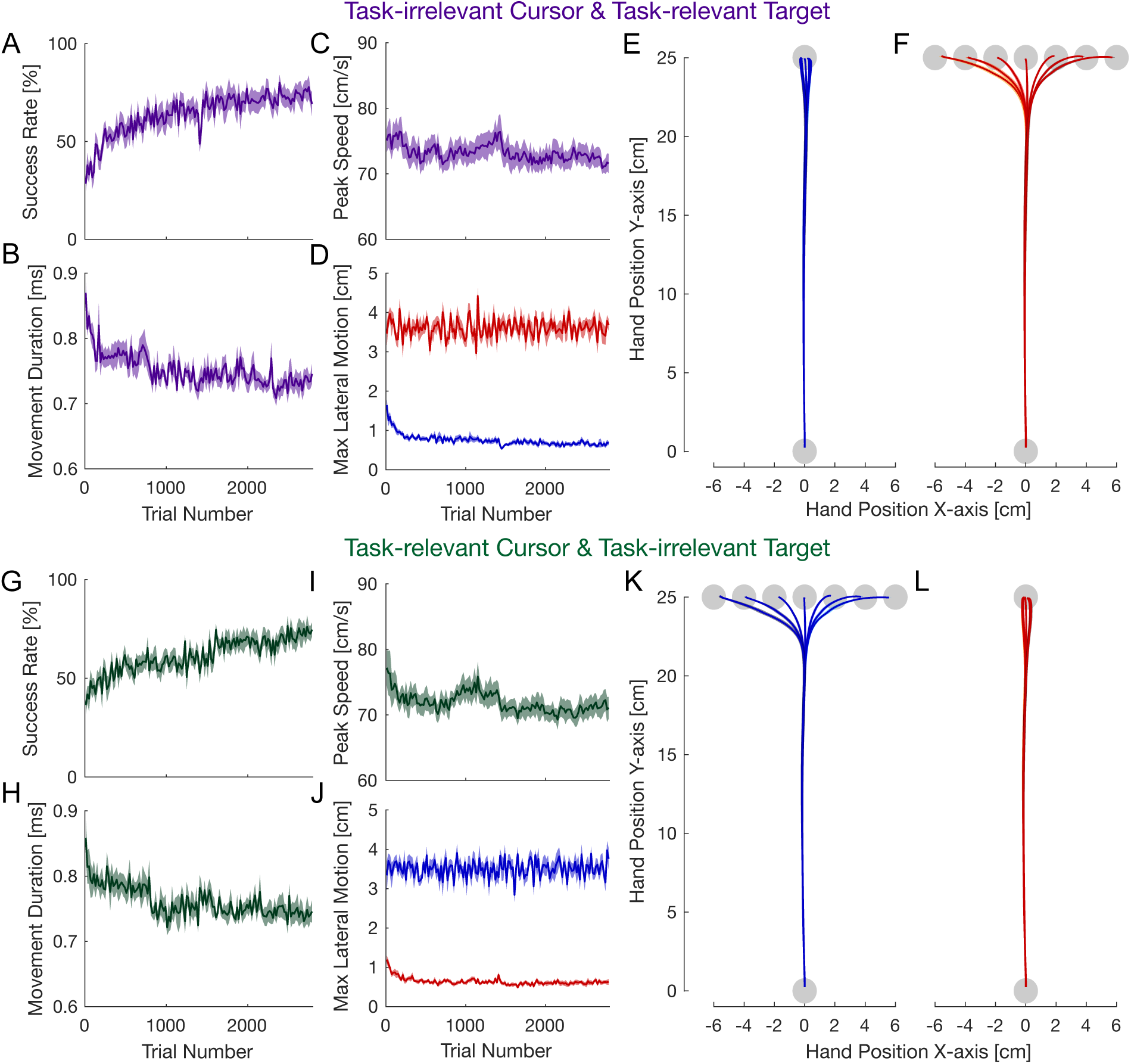
Performance in the two experimental conditions. **A-F**: Task-irrelevant cursor and task-relevant target condition. **A**. Mean success rate over the 2800 trials. Shaded region indicates standard error of the mean. **B**. Movement duration. **C**. Peak forward speed. **D**. Average maximum absolute lateral motion on the non-probe trials, with trials in which the cursor moved shown in blue and trials in which the target moved shown in red. **E.** Hand trajectories during the trials with task-irrelevant cursor motion. The mean (solid line) and standard error of the mean (shaded region) are shown for each of the seven possible types of cursor motion. Trajectories are shown only for the second session (trials 1401-2800). **F**. Hand trajectories during the trials with task-relevant target motion. As the target moves during the trial, the final position of the target for each of the seven different motions of the target are shown. **G-L**: Task-relevant cursor and task-irrelevant target condition. **G**. Mean success rate. **H**. Movement duration. **I**. Peak forward speed. **J**. Average maximum absolute lateral motion on the non-probe trials. **K.** Hand trajectories during the trials on the second session (trials 1401-2800) with task-relevant cursor motion. As the cursor shift remains at the end of the movement, the relative final position of the target for each of the seven different motions is shown. **L**. Hand trajectories during the trials with task-irrelevant target motion.

Participants showed similar performance and adaptation in the other environment in which the cursor motion was task-relevant and the target motion task-irrelevant. Again, success rate increased (t_9_=14.85; p<0.0001) from 42.3% (first 100 trials) to 73.5% (last 100 trials) (Fig 1G) and movement duration decreased (t_9_=5.08; p=0.0007) over the two sessions (Fig 1H). However, there was no significant change in peak forward speed (t_9_=2.02; p=0.0745) over the course of the experiment (Fig 1I). The lateral motion to cursor and target motion was now reversed (Fig 1J). Participants now made straight trajectories to the target when the target underwent the task-irrelevant motion (Fig 1F). However, the hand trajectories during the cursor motion were now strongly corrected, moving such that the final hand position matched the needed offset for the cursor to end within the target location (Fig 1E). Overall, it appears that participants were able to learn to control the hand trajectories appropriately in both environments, correcting when the motion was task-relevant and needed to be corrected, and ignoring the motion when it was task-irrelevant.

To characterise the changes to cursor and target motion in the two conditions, we investigated the lateral hand acceleration for each of the different motion patterns. In the task-irrelevant cursor and task-relevant target motion condition (Fig 2), the time course of the cursor motion and target motion underwent similar initial motion, but the cursor was returned to the hand position (Fig 2A), whereas the target motion needed to be corrected for (Fig 2C). The lateral hand acceleration during the cursor motion (Fig 2B) occurs in the direction as if to correct for the cursor motion, but the response remains small and is rapidly corrected in the opposite direction (around 300 ms after the start of the cursor motion). In contrast, the hand acceleration during target motion has a similar onset time but increases and remains large over the subsequent 400 ms (Fig 2D). To examine the relative hand acceleration in visuomotor feedback responses latencies, we quantified these measures over both an early and late time window. While the acceleration produced in response to target motion appeared slightly larger in the early window than that to cursor motion (Fig 2E,G), this difference was enhanced in the later time window (Fig 2F,G). These results already suggest that the visuomotor responses to cursor and target motion might be downregulated and unregulated respectively, but this would need to be confirmed by examine these responses in the second environmental condition, where the task-relevancy is switched for cursor and target motion.

**Figure 2.**
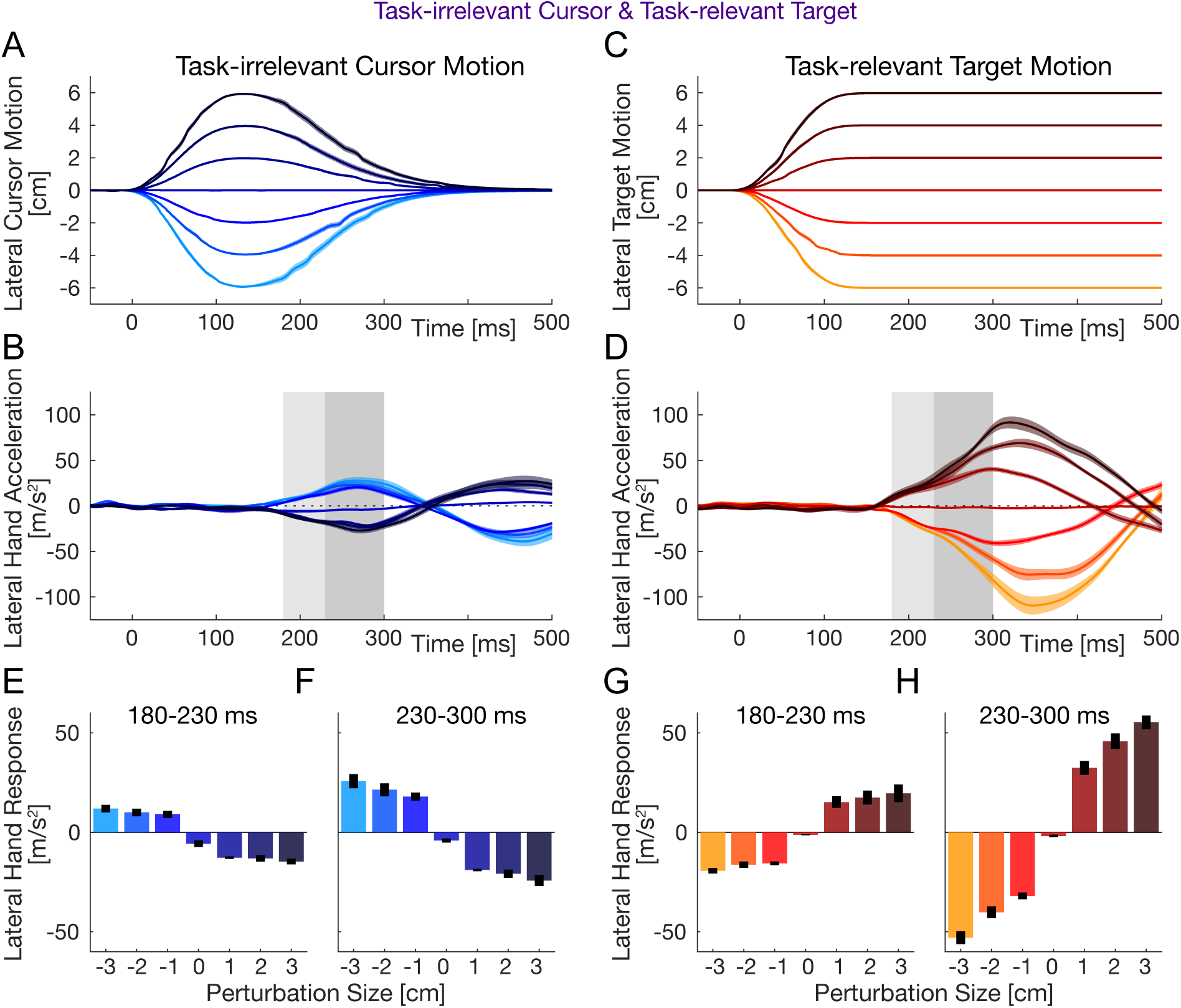
Kinematic correction to task-irrelevant cursor motion and task-relevant target motion. **A.** The lateral cursor motion which was applied after the participant had moved 7.5 cm from the start position plotted as a function of time from the start of this motion. Each colour denotes one of the seven different amplitudes of motion. Time of 0 indicates onset of the lateral visual motion of cursor. **B.** The lateral hand acceleration measured in response to each of the seven different task-irrelevant cursor motions. The color corresponds to the perturbation magnitude. The light grey region indicates the early involuntary time window (180-230 ms) whereas the darker grey region indicates the late time window (230-300ms). **C.** The lateral target motion that was applied after the participant had moved 7.5 cm from the start posittion plotted as a function of time. **D.** The lateral hand acceleration measured in response to each of the seven different task-relevant target motions. **E.** The hand response in the early time window to the shift in the cursor motion. Each bar represents the mean lateral acceleration in the early time window for the seven amplitudes. Error bars indicate the standard error of the mean. **F.** The hand response in the late time window to the shift in the cursor motion. **G.** The hand response in the early time window to the shift in the target motion. **H.** The hand response in the late time window to the shift in the target motion.

By examining the hand acceleration responses in the task-relevant cursor and task-irrelevant target motion condition (Fig 3), we can clearly see that the gains of the feedback responses are now switched, with higher responses to the cursor motion and lower responses to the target motion. The corrective lateral hand acceleration is large in response to the maintained cursor motion (Fig 3B), but small and switches direction in response to the task-irrelevant target perturbations (Fig 3D). Again, by examining these measures in the early and late time windows, we can see that the relative gain of the two measures in now reversed, with slightly large responses to cursor motion compared to target motion in the early window (Fig 3E,G) and large different in the late window (Fig 3F,H). These results provide some initial evidence that in the two different visual environments, the participants were able to switch the gain of the visuomotor feedback responses to cursor and target motion differentially.

**Figure 3.**
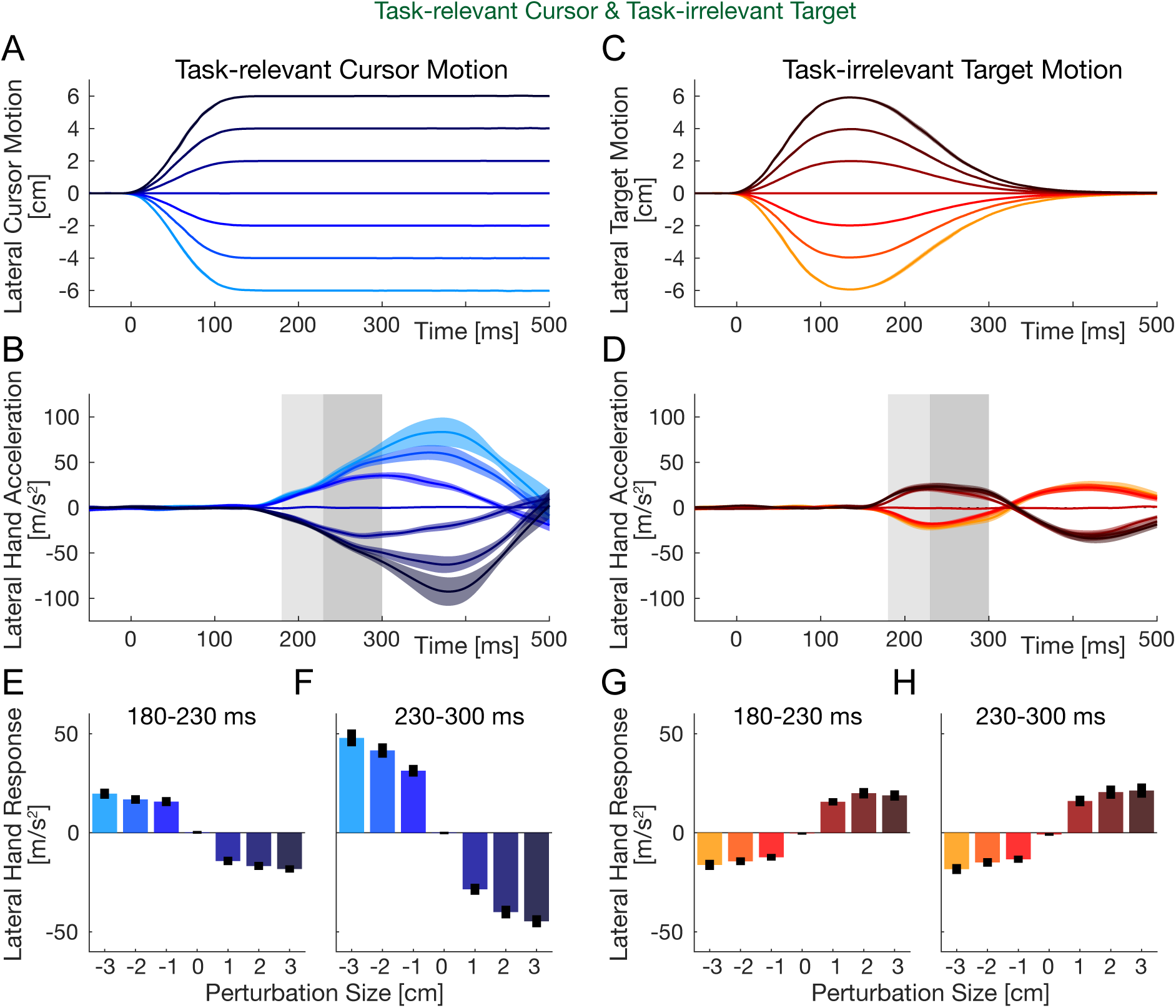
Kinematic corection to task-relevant cursor motion and task-irrelevant target motion. **A.** The lateral cursor motion which was applied after the participant had moved 7.5 cm from the start posittion plotted as a function of time. Each color denotes one of the seven different amplitudes of motion. **B.** The lateral hand acceleration measured in response to each of the seven differnet cursor motions. The color corresponds to the pertubation magnitude. The light grey region indicates the early involuntary time window (180-230 ms) whereas the darker grey region indicates the late time window (230-300ms). **C.** The lateral target motion that was applied after the participant had moved 7.5 cm from the start posittion plotted as a function of time. **D.** The lateral hand acceleration measured in response to each of the seven different target motions. **E.** The hand response in the early time window to the shift in the cursor motion. Each bar represents the mean lateral acceleration in the early time window for the seven amplitudes. Error bars indicate the standard error of the mean. **F.** The hand response in the late time window to the shift in the cursor motion. **G.** The hand response in the early time window to the shift in the target motion. **H.** The hand response in the late time window to the shift in the target motion.

### Visuomotor Responses

The kinematic responses to the two different environments appeared to be modified according to the task-relevancy of the cursor or target motion (Figures 2 and 3). However, the later visual motion of the cursor and target were also different in the two experimental conditions, which might limit our ability to directly compare. This is further complicated by the slow, smooth onset of the visual motion which makes determining the exact onset time difficult for quantifying specific time windows. To directly test whether the gain of the visuomotor feedback responses could be independently controlled, here we directly assessed the visuomotor responses using identical probe trials in both environments. Throughout the entire experiment, the visuomotor feedback responses were measured using probe trials on random trials in which the hand was constrained to a straight-line movement and the force response to the visual stimuli was measured against the channel wall (Methods). The force responses were measured in response to seven different sizes of probe trials.

As the gain of visuomotor responses can adapt to the environmental conditions with training ^22,24,42,43^, we first examined the changes in the visuomotor feedback responses over both the early (180-230 ms) and late (230-300 ms) time windows throughout the experimental condition (Fig 4). In the task-irrelevant cursor and task-relelevant target motion condition, the visuomotor feedback responses to cursor perturbations (probe trials) quickly decreases over the first 5 blocks of trials and is fairly low for the subsequent 35 blocks in both the early and late time windows (Fig 4A,B). In contrast the visuomotor feedback responses to target perturbations in both time intervals remains high over the course of the full experiment. In the opposite condition (task-relevant cursor and task-irrelevant target motion), the opposite changes in feedback responses are seen (Fig 4C,D). The visuomotor response to the cursor perturbations remains high throughout the experiment, whereas the visuomotor feedback responses to target motion reduce rapidly over the first five blocks and then remain low throughout the experiment.

**Figure 4.**
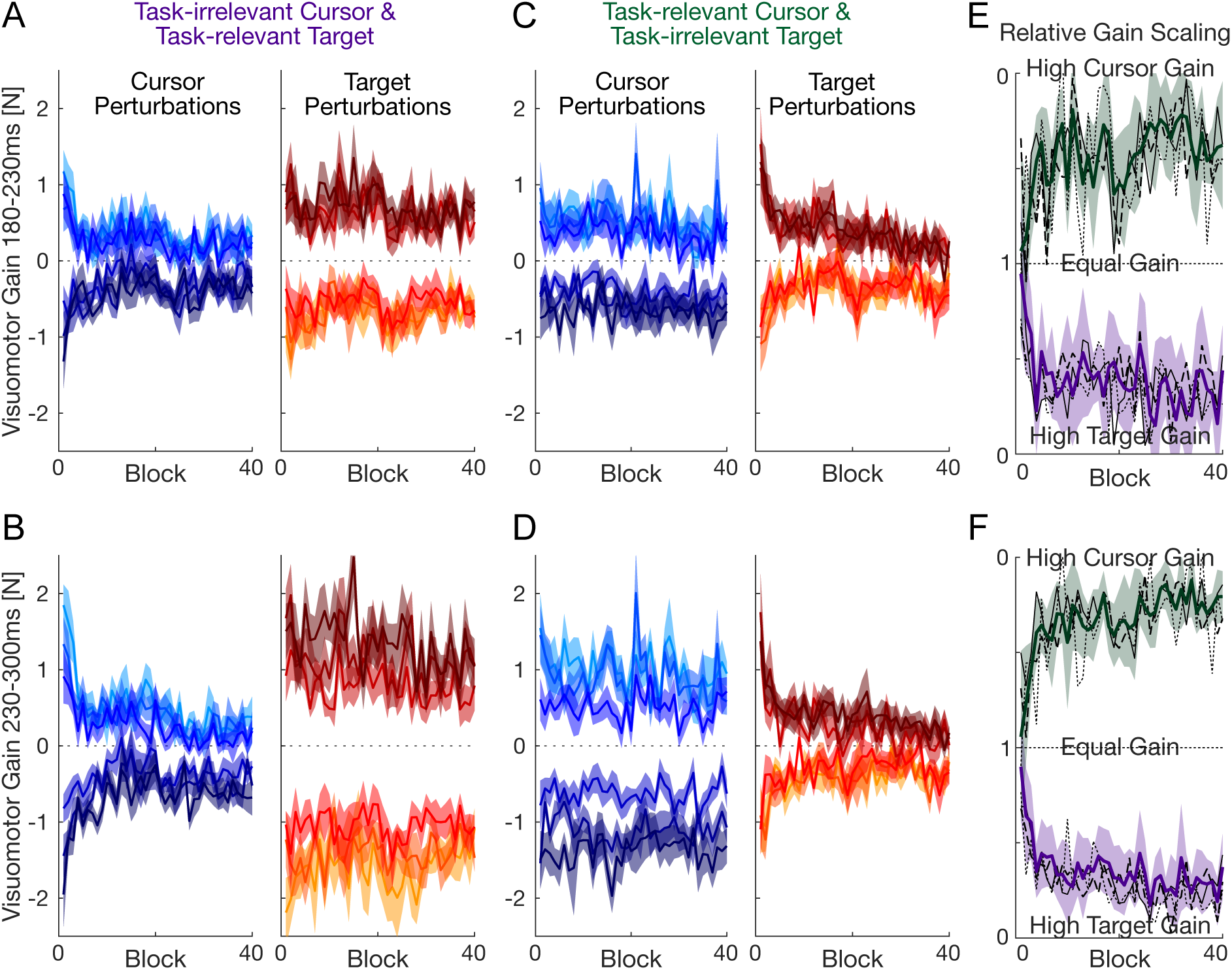
Time course of adaptation of visuomotor feedback responses. **A.** Early visuomotor feedback gains in protocol 1 (task-irrelevant cursor and task-relevant target motion). Responses are shown to cursor probe trials (blue) and target probe trials (red) over each block of the experiment. The visuomotor response to each perturbation size was quanitifed between 180-230 ms. Colors indicate the size of the visual perturbation of the cursor from −3 cm (light blue) to + 3 cm (dark blue) or of the target from −3 cm (light orange) to +3 cm (dark red). **B.** Late visuomotor feedback gains (230-300ms) in protocol 1 (task-irrelevant cursor and task-relevant target motion). **C.** Early visuomotor feedback gains in protocol 2 (task-relevant cursor and task-irrelevant target motion). **D.** Late visuomotor feedback gains (230-300ms) in protocol 2. **E.** Relative gain scaling between the cursor and target responses for each block for both experimental conditions (green and purple) for the early time window (180-230 ms). Gain scaling was calculated as task-irrelevant divided by task-relevant for each condition and ploted between 0 and 1 for both enviornments. On the top values less than one indicate larger cursor feedback responses and on the bottom values less than one indicate larger target feedback responses. Error bars are 95% confidence intervals. Black lines indicate mean values across participants for the 1 cm (dotted), 2cm (dashed) and 3 cm (solid) size perturbations. **F.** Relative gain scaling between the cursor and target responses for each block for both experimental conditions (green and purple) for the late time window (230-300 ms).

In order to quantify the changes in the feedback gains between the cursor and target perturbations we calculated the ratio of visuomotor feedback responses block by block in both environments as the task-relevant feedback gain divided by the task-irrelevant feedback gain. We can then directly compare the scaling of the target and cursor feedback gains in the two experimental conditions. In the early involuntary time window, while the ratio is close to 1 in both environments at the start of the experiment, with overlapping 95% confidence intervals, this rapidly decreases, and a clear difference in the scaling of the target and cursor feedback gains is seen in both environments (Fig 4E). Although the ratio of visuomotor feedback gains is fairly variable in this early time window, the ratio remains clearly different from 1 (based on the 95% confidence intervals) and more critically is significantly different in the two environments (no overlap in the two confidence intervals after the first few blocks). This difference in the ratio between target and cursor feedback gains is even more apparent in the later time window (Fig 4F) which again rapidly decreases from equal scaling in the first block to around 0.3 by the end of the experiment in both conditions. Again, the scaling is significantly different from 1, and is significantly different in the two experimental conditions. Overall, we find significant evidence for independent scaling of visuomotor responses to target and cursor motion which develops as participants experience to two conditions.

The results of the block-by-block analysis (Fig 4) demonstrated that most of the rapid changes in the visuomotor feedback responses were occurred wihin approximately the first 5 or so blocks in each condition. We therefore examined the final level of visuomotor feedback gains after the initial 10 blocks of trials (after the first 700 trials in each condition) using the final 30 blocks of trials. In the task-irrelevant cursor and task-releveant target condition (Fig 5), we find that the visuomotor feedback responses are smaller for cursor perturbations than target perturbations (Fig 5C,D), and that this effect is clear across both the early and late time windows (Fig 5 E-H). By examining the relation scaling between the cursor and target visuomotor responses, we find a clear difference from 1 (Fig 5I,J), showing that the visuomotor feedback responses are larger for target than cursor perturbations. Importantly, this is not only for the mean across participants; every one of the ten participants is significantly less than 1.0 for both the early and late time windows, with scaling generally below 0.5. A t-test confirmed that the scaling ratio is less than 1.0 for both the early (t_9_=12.6; p<0.0001) and late (t_9_=22.3; p<0.0001) time windows.

**Figure 5.**
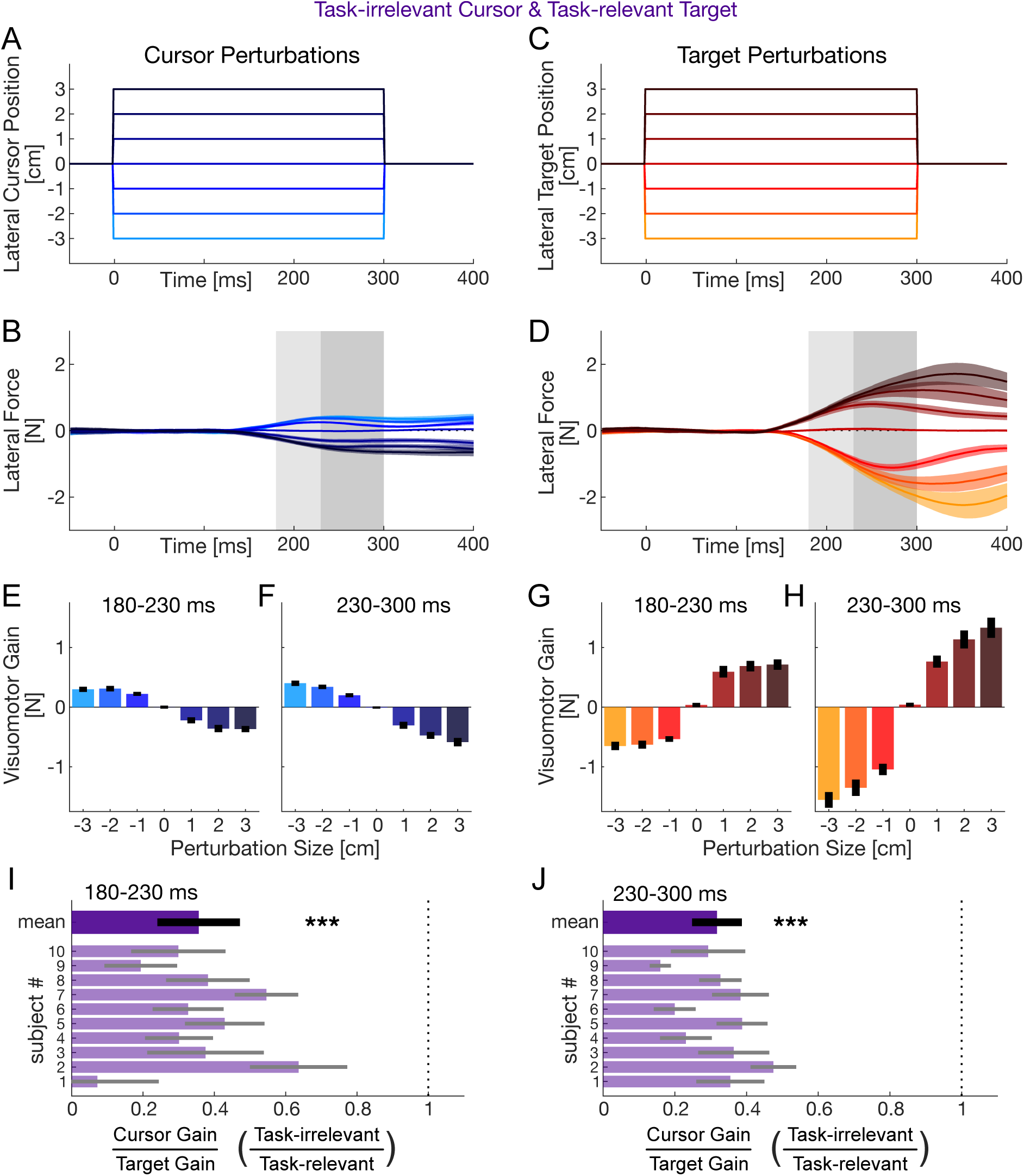
Tuning of the visuomotor feedback repsonses in protocol 1 (task-irrelevant cursor and task-relevant target motion) shows downregulation of the feedback responses to the cursor compared to the target. **A.** The cursor motion on the probe trials. **B.** The lateral force response to the cursor motion. Colors match the perturbation amplitudes in A. Colored shaded region indicates the standard error of the mean across participants. Light grey and dark grey bars show the region over which the early and late feedback responses were quantified. **C.** The target motion on the probe trials. **D.** The lateral force response to the target motion. **E.** Early visumotor feedback responses to cursor motion (mean and standard error of the mean from 180-230 ms). **F.** Late visuomotor feedback responses to cursor motion. **G.** Early visumotor feedback responese to target motion. **H.** Late visuomotor feedback responses to target motion. **I.** Scaling of cursor gain to target gain (magnitude of task-irrelevant feedback responses with respect to task-relevant feedback repsonses) during the early feedback window. Under this protocol every participant showed a reduced cursor response compared to target response. The dotted line at 1.0 indicates equal sized responses. At the top the mean and 95% confidence intervals of the mean across participants is illustrated. Individual participant means are shown with 95% confidence intervals indicated by the grey bars. **J.** Scaling of cursor gain to target gain (magnitude of task-irrelevant feedback responses with respect to task-relevant feedback repsonses) during the late feedback window.

In the final 30 blocks of trials in the task-irrelevant cursor and task-releveant target condition (Fig 6), we see the opposite effect. There are larger visuomotor feedback responses for cursor perturbations than target perturbations (Fig 6C,D), across both the early and late time windows (Fig 6 E-H). By examining the relation scaling between the cursor and target visuomotor responses, we again see a clear difference from 1 (Fig 6I,J), showing that the visuomotor feedback responses are now larger for cursor perturbations than target perturbations. Moreover, this is again seen for every participant, with scaling values significantly less than 1.0 for both the early and late time windows. A t-test confirmed again that the scaling ratio is less than 1.0 for both the early (t_9_=17.4; p<0.0001) and late (t_9_=22.6; p<0.0001) time windows. Overall we find that the gain of the visuomotor feedback responses can be independently tuned for visual feedback of the cursor and the target depending on the visual environment.

**Figure 6.**
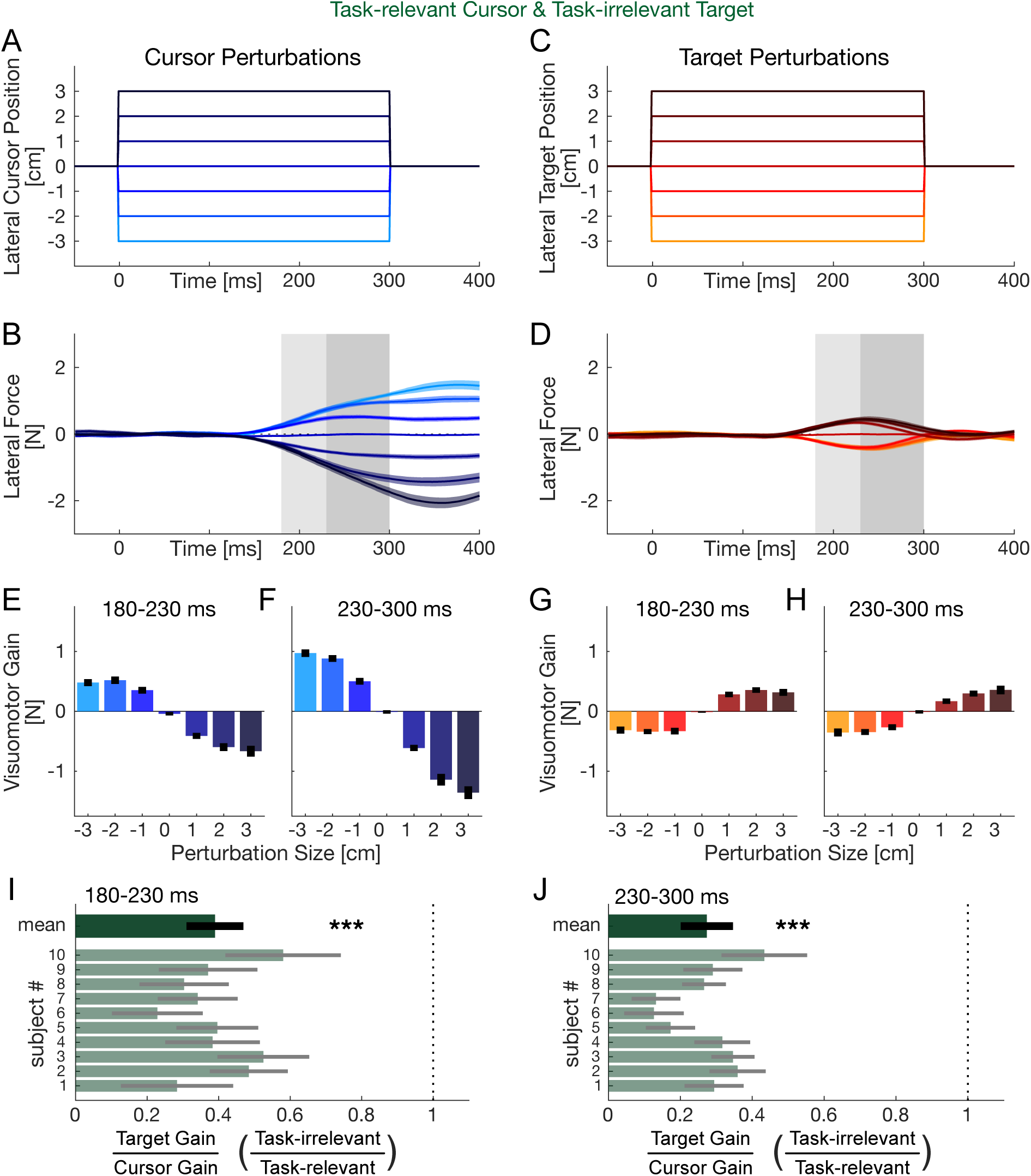
Tuning of the visuomotor feedback repsonses in protocol 2 (task-relevant cursor and task-irrelevant target motion) shows upregulation of the feedback responses to the cursor compared to the target. **A.** The cursor motion on the probe trials. **B.** The lateral force response to the cursor motion. Colors match the perturbation amplitudes in A. Colored shaded region indicates the standard error of the mean across participants. Light grey and dark grey bars show the region over which the early and late feedback responses were quantified. **C.** The target motion on the probe trials. **D.** The lateral force response to the target motion. **E.** Early visumotor feedback responses to cursor motion (mean and standard error of the mean from 180-230 ms). **F.** Late visuomotor feedback responses to cursor motion. **G.** Early visumotor feedback responese to target motion. **H.** Late visuomotor feedback responses to target motion. **I.** Scaling of target gain to cursor gain (magnitude of task-irrelevant feedback responses with respect to task-relevant feedback repsonses) during the early feedback window. Under this protocol every participant showed a reduced target response compared to cursor response. The dotted line at 1.0 indicates equal sized responses. Mean and 95% confidence intervals are shown for individual participants and for the mean across participants. **J.** Scaling of target gain to cursor gain (magnitude of task-irrelevant feedback responses with respect to task-relevant feedback repsonses) during the late feedback window.

## Discussion

Participants completed four days of reaching experiments designed to test whether the rapid visuomotor feedback responses to cursor and target motion can be tuned independently and simultaneously. Participants reached in two visual environments: in one the cursor motion was task-irrelevant while the target motion was task-relevant, and in the other these roles were reversed. In both environments, participants learned to correct for the task-relevant displacements while largely ignoring the task-irrelevant ones, improving their success rate and shortening their movement duration over training. Crucially, we probed the feedback responses throughout both environments using identical lateral perturbations of the cursor or the target. Participants upregulated the visuomotor response to whichever signal was task-relevant while simultaneously downregulating the response to the task-irrelevant signal, and were able to reverse this pattern when the task relevance of the two signals was switched. This differential tuning developed rapidly, within the first five blocks of each environment, was already present in the early, involuntary interval (180–230 ms), and strengthened in the later interval (230–300 ms). The effect was highly consistent: every participant showed a task-dependent gain ratio well below one in both intervals and both environments. Together, these results demonstrate that the gains of the visuomotor feedback responses to hand and target information can be set independently of one another, at the same time, as a function of task-relevance.

Our findings extend a line of work showing that visuomotor feedback gains are not fixed but are continuously tuned to the demands of the task. Franklin and Wolpert^22^ demonstrated that the involuntary response to a shift in the visual hand position is upregulated when such discrepancies are task-relevant, and a downregulation occurs when the discrepancies are task-irrelevant i.e., when they require no correction. This tuning was subsequently shown to be highly flexible. For example, feedback gains can be set according to different limb states and for different perturbation directions within a single posture, such that responses on one side of the movement are upregulated while responses on the other side are suppressed ^28^. The same scheme applies to how visual feedback of the hand is used during pointing: participants flexibly adjust the feedback control laws that map sensory signals onto motor commands according to the accuracy demands of the task, correcting along behaviourally relevant dimensions while discounting those that do not affect task success ^26^. Here, we partitioned task relevance across two distinct visual signals (cursor and target) that were displaced within the same reaching movements. Participants assigned opposite gains to these two channels at the same time, and appropriately swapped that assignment when the environment changed. To our knowledge, this is the first demonstration that the hand-related and target-related visuomotor responses can be independently and oppositely tuned within a single behaviour.

Our results help disambiguate competing accounts of how visual information guides reaching. The classical difference-vector view proposes that the brain extracts the positions of the hand and the target, computes a single difference vector between them, and uses this combined error signal to drive corrective motor commands ^41^. Such a scheme ultimately implies the presence of a single feedback channel and therefore predicts that the responses to hand and target displacement are intertwined. However, Reichenbach and colleagues ^39^ reported that visual attention facilitates the processing of target information but not of hand information, pointing to a dedicated visuomotor binding mechanism that grants reafferent hand information preferential and independent access to motor control. Moreover, the difference-vector model was tested directly by measuring responses across a large matrix of combined cursor and target displacements ^40^; they found that the early responses behaved as two independent contributions that were only later combined into an accurate difference vector. Our results add decisively to this line of evidence: because the cursor and target responses can be assigned opposite gains at the same time, they cannot be read out from a single shared error signal, at least over the intervals we examined. The most parsimonious interpretation is that hand and target information are handled by separable feedback controllers whose gains can be set independently, in line with the independent processing found to dominate the early response ^40^. This arrangement is also consistent with the parallel specification of feedback gains observed when the two hands pursue independent goals ^44^.

The differential gains demonstrated in our study did not require extensive training. Most of the change occurred within the first few blocks of each environment and was then maintained for the remainder of the two sessions. This rapid, experience-dependent reweighting is consistent with the framework of optimal feedback control, in which feedback gains are adjusted so that corrective effort is spent on task-relevant deviations while task-irrelevant fluctuations are left uncorrected ^37,38,45^. This is the minimal intervention principle in action: corrective gain is allocated only where deviations jeopardise the goal. The speed with which our participants reweighted the two channels also fits earlier evidence that visuomotor feedback gains are not slowly re-learned but updated rapidly, on the timescale of a few movements, as task demands change ^43^. The finding that the appropriate gains were already in place from the earliest involuntary interval further suggests that experience reconfigures the feedback controller itself, presetting channel-specific gains, rather than merely adding later voluntary corrections. Tuning the cursor and target channels separately is precisely what such a controller should do when only one of the two signals carries information relevant to the goal. The temporal profile of the effect is also informative. Independent tuning was present in the early involuntary interval but became more pronounced later in the response. This aligns with evidence that the visuomotor feedback response unfolds in stages, with an initial rapid approximation that is progressively refined ^40,46^, and suggests that the capacity for independent gain control is available from the earliest, largely involuntary phase of the response and is then further shaped by slower processes.

A natural question is how two functionally independent feedback systems might be implemented in the nervous system. Rapid visuomotor responses have long been linked to a subcortical pathway through the superior colliculus, a phylogenetically conserved midbrain area that receives converging input from multiple sensory modalities (visual, auditory and somatosensory)^47^. Via tecto-reticulo-spinal projections, it can drive rapid muscle responses to visual stimuli at latencies of roughly 80–120 ms ^12,48^. These rapid responses are tuned toward the location of the visual stimulus and are largely automatic, yet they are not rigid: they can be enhanced or suppressed by top-down, cortically derived task context ^48,49^. Recent work using a jumping-target task likewise argues that online corrections are not a separate class of movement but instead reflect a nested control system, comprising an early subcortical phase (∼80–120 ms) followed by a cortical phase (>130 ms), both of which can be preset by task demands ^50^ (delays to production of electromyographic activity). Although the voluntary driven response to produce force responses is thought to be around 230 ms ^22,46^, the electromechanical delay of arm muscles is only around 40-50 ms ^51,52^, arguing for potential cortical involvement starting around 180 ms. This means that our measurements could include substantial cortical contributions, perhaps exploiting the functional division between dorsal action-related and ventral perception-related visual processing ^53^.

On the cortical side, the posterior parietal cortex is a strong candidate node for the longer-latency component. Disrupting it with transcranial magnetic stimulation selectively abolishes the online corrections evoked by target jumps while sparing reaches to stationary targets ^54^, consistent with a forward-model architecture in which an internal estimate of hand state is continuously compared against the goal to drive fast corrections ^55^. Moreover, the parietal cortex has been proposed to maintain separable estimates of bodily and environmental states ^56^, an organisation well suited to supporting the independence between hand- and target-related control that we observe, with the two estimates converging onto shared motor output only downstream.

Several limitations pertain to our main conclusion of an independence between hand and target visuomotor feedback control. While our main finding was present in every participant and across both the early and late response intervals, we probed only lateral displacements applied at a single point in the movement; it remains to be established whether independent tuning generalises across perturbation timing, across amplitudes beyond the tested range, and across movement directions. In addition, behavioural measurements cannot identify the neural locus of the two controllers or determine whether they partly share intermediate processing stages.

Combining this paradigm with electromyography to dissociate the express and longer-latency components ^46,50^, or with neuroimaging and the perturbation of candidate neural structures would help clarify whether the independence we observe reflects truly parallel pathways, partially independent pathways or a single pathway with signal-specific gating.

We show that rapid visuomotor feedback responses to cursor and target motion can be tuned independently, and in opposite directions, within a single visual environment according to the task relevance of each signal. This independent and simultaneous control adds direct functional evidence that hand and target information are handled by separate feedback systems whose gains are flexibly set to facilitate the task at hand.

## Methods

### Experimental Participants

Ten neurologically healthy young adults participated in this series of experiments (6 males and 4 females: aged 23.6 ± 2.4, mean ± SD). All participants were naïve to the purpose of the study and right-handed according to the Edinburgh handedness inventory ^57^. The study was approved by the Ethics Committee of the Medical Faculty of the Technical University of Munich. All participants provided a written informed consent before participating in the study. Each participant made movement under two different protocols which were designed to test whether the visuomotor feedback response could be independently modulated to perturbations of the cursor and the target. Half of the participants started the experiment with protocol 1 followed by protocol 2, whereas the other half of the participants performed protocol 2 followed by protocol 1. Each protocol was performed for two sessions.

### Apparatus and Setup

Participants performed reaching movements to a target while grasping a robotic manipulandum (Fig. 7A). Participants were seated in an adjustable chair in front of the robotic rig with their shoulders restrained by a harness. The participants’ right arm rested on an air sled and they grasped the handle of a vBOT robotic interface ^58^ with the right hand (Fig. 7A). A six-axis force transducer (ATI Nano 25; ATI Industrial Automation) measured the end-point forces applied by the participant at the handle. The position of the vBOT handle was calculated from joint-position sensors (58SA; Industrial Encoders Direct) on the motor axes. Position and force data were sampled at 1 kHz. Visual feedback was provided using a computer monitor mounted above the vBOT and projected into the plane of the movement via a mirror. The participants hand and arm were hidden by the virtual reality system, preventing direct visual information of their hand position. The exact onset time of any visual stimulus presented to the participant was determined using the video card refresh signal, which had been confirmed with an optical signal.

**Figure 7.**
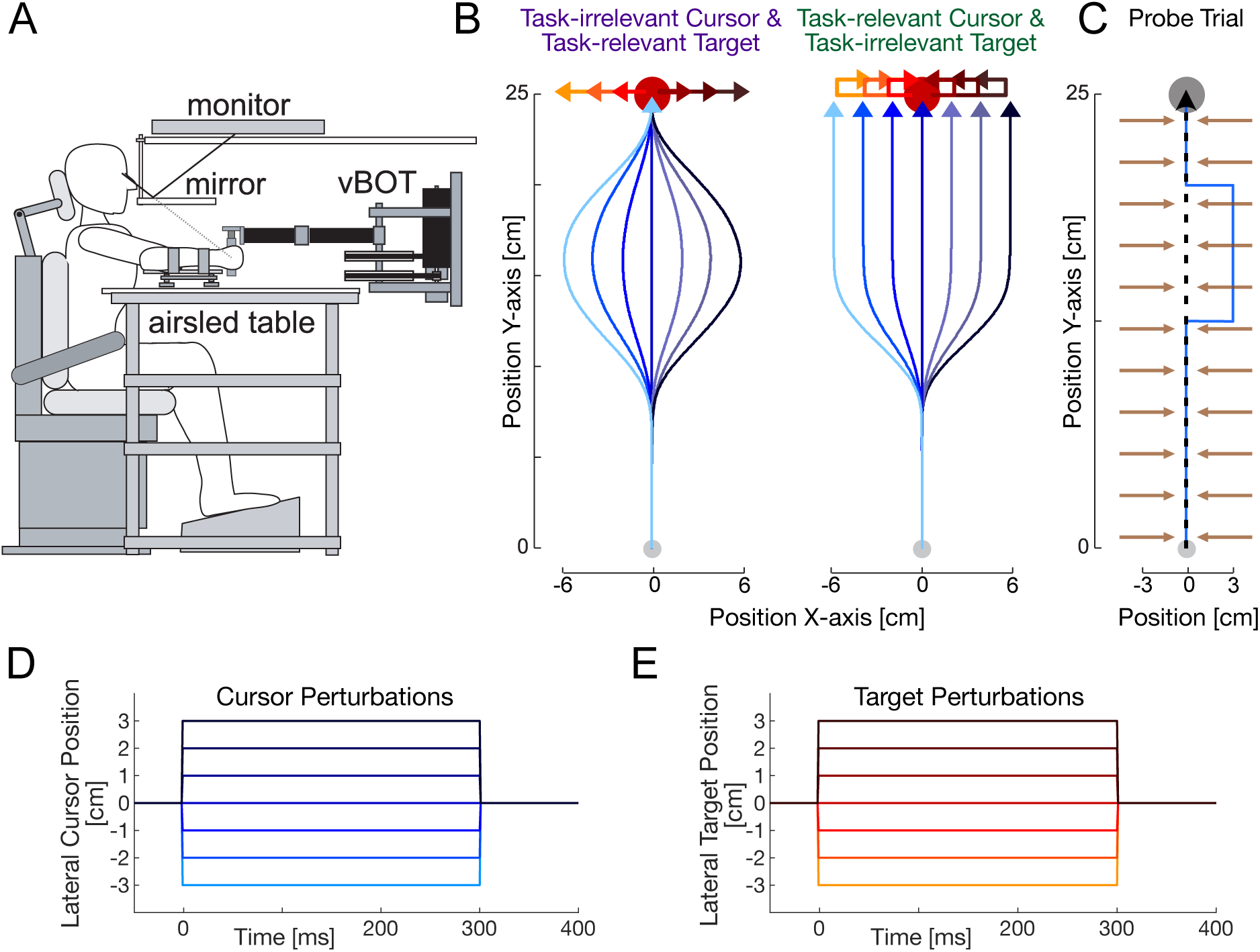
Experimental Design. **A**. Participants performed reaching movements while grasping the vBOT robotic manipulandum. **B**. Each participant performed reaching movements in two different visual environments. In the task-irrelevant cursor and task-relevant target condition, the target underwent smooth motion with an offset at the end of the movement that needed to be corrected for (red) whereas the cursor underwent smooth motion with no final offset (blue). In the task-irrelevant cursor and task-relevant target condition, these movements were reversed. **C**. Random probe trials were interspersed in both environments to independently measure the visuomotor feedback responses. On these trials the hand motion (dotted black line) was constrained to a straight movement to the target by a virtual mechanical channel (brown arrows) while the visual representation of the cursor (blue line) was perturbed laterally. **D**. Seven different sizes of cursor perturbations. **E**. Seven different sizes of target perturbations.

### Trial Procedure

Individual participants completed 4 sessions, each 1400 trials long, on 4 days. In total, each participant completed 5600 trials. All trials were self-paced; a yellow 0.5 cm diameter cursor was projected to instantaneously represent their hand position. Movements were made from a 0.8cm diameter start circle (grey circle which became white once participants had moved the cursor into the start circle) to a 0.5 cm diameter yellow target circle, both of which were centred in front of the participant. Participants initiated a trial by moving the hand cursor into the start circle and holding it for 1.1 s. A beep then indicated that the participant could begin the movement to the target. The duration of the movement was determined from the time that the participant’s hand exited the start target until the time that the participant’s hand entered the final target. The distance between the centres of the start and target circles was 25.0 cm. Successful movements were defined as those that entered the target without overshooting and with movement durations in the range 700 ± 100 ms and were accompanied with feedback (“good” or “great”). “Great” was presented when the movement was in the range, 700 ± 50 ms, whereas “good” was presented otherwise (600-650 ms or 750-800 ms). When participants performed successful movements, a counter increased, and participants were instructed to increase the number of the counter as much as possible throughout the experiment. When participants performed unsuccessful movements, they were provided with feedback as to why the movement was not considered successful such as “too fast” (duration <600 ms), “too slow” (duration >800 ms) or “overshot target” (if the movement exceeded the final target by 1.0 cm). Participants were then free to return to the start point to initiate the next trial while the feedback was provided about the success of the previous trial. The visual feedback about the location of the hand (cursor) was only provided during the return movement once the hand was within 5 cm of the start target.

### Probe trials

Throughout the entire experiments, visuomotor feedback responses were examined with visual perturbations of the cursor or target (Fig 7C) as we have done in previous studies ^24,27,40,42,43^. In the middle of the movements to the target, visual perturbations of either the cursor or the target were performed laterally by one of seven possible distances [−3, −2, −1, 0, 1, 2, 3] cm when the cursor reached the middle of the movement (12.5 cm from the start). Together, this resulted in 14 different visual perturbations. During cursor perturbations, the cursor representing the hand position was laterally jumped away from the current hand position, held at a fixed distance from the actual hand trajectory for 300 ms, and then returned to the actual hand position for the rest of the movement (Fig 7D). During target perturbations, the target was laterally jumped, held at the new location for 300 ms and then returned to the original location for the rest of the movement (Fig 7E). During such probe trials, the hand was physically constrained to the straight path between the start and final targets using a mechanical channel trial ^22,59^ generated by the vBOT. The mechanical channel was implemented as a stiffness of 6,000 N/m and damping of 2 N·m^−1^·s^−1^ for any movement lateral to the straight line joining the starting location and the middle of the target. This constrains the physical hand location using the channel such that no lateral change in the arm configuration occurs. The force produced in response to the visual perturbation can be measured against the channel wall with the force sensor to measure the gain of the visuomotor feedback response ^22,23,28,39,40^. As the visual perturbation returns to the actual hand trajectory, participants do not need to respond to the visual perturbation to produce a successful movement to the target. These probe trials were randomly applied during movements in a blocked fashion such that one of each perturbation type was applied within each block of trials.

### Protocol

To determine whether these visuomotor reflex gains to the cursor and target perturbations could be modulated independently and simultaneously, participants were exposed to different distributions of sensory discrepancies to the cursor and to the target. In one experimental condition the cursor was exposed to task relevant discrepancies and the target to task irrelevant discrepancies, while during the other condition this was reversed. Under both conditions, the visuomotor feedback responses were probed independently using identical probe trials.

#### Experimental Condition 1 (task-irrelevant cursor and task-relevant target)

In this protocol, during reaching movements either the visual cursor representing the hand or the visual target representing the final target were smoothly moved laterally (Fig 7B). During some of the trials the visual target moved laterally to 1 of 7 amplitudes (including 0) and remained at this location for the rest of the movement. In contrast, while the visual cursor underwent the same initial movement to one of seven amplitudes (including 0), it was then smoothly moved back to the original location (matching the hand position) before the end of the movement. Specifically, after the participant reached the point in the trajectory that was 30% of the distance to the target (7.5 cm from the start), the visual target underwent a smooth (minimum jerk ^60^) movement laterally to the movement direction to a distance from the set [−3, −2, −1, 0, 1, 2, 3] cm in the subsequent 30% of the movement, remaining at this position laterally for the rest of the movement. Participants were required to produce the appropriate response to bring the hand cursor into the target and be credited with a successful trial, which is why this visual disturbance is termed a task-relevant discrepancy ^22,28^. On other trials, once the participant reached 30% of the distance to the target, the visual cursor representing the hand position moved laterally with a minimum jerk trajectory to one of seven distances [−3, −2, −1, 0, 1, 2, 3] cm relative to the hand location in the subsequent 30% of the movement, before moving back to match the hand position with a minimum jerk trajectory over the next 30% of the movement. It then remained matching the hand position for the final 10% of the movement. In this condition, while the lateral change in the visual location of the hand position produces a visuomotor discrepancy, the participants should ignore both the size and direction of these sensory discrepancies as they do not need to be compensated for to successfully produce the movement to the target. These are termed task-irrelevant sensory discrepancies ^22,28^. All sensory discrepancy trials were performed in the null field such that participants could easily correct their hand motion as desired. Under these conditions, participants should ideally correct for target perturbations but ignore cursor perturbations if they can control these feedback responses independently.

#### Experimental Condition 2 (task-relevant cursor and task-irrelevant target)

In this protocol the reversed conditions were applied to the cursor and targets: task-relevant sensory discrepancies were applied to the cursor and task-irrelevant sensory discrepancies were applied to the target (Fig 7B). Again, at 30% of the distance to the target, the visual cursor smoothly (minimum jerk) moved away from the hand trajectory to 1 of 7 amplitudes (including 0) in the subsequent 7.5 cm and remained at this location with respect to the hand position for the rest of the trial. Here participants need to adjust their trajectory to size and direction of the visual discrepancy in order to reach the target. On the other trials, after participants were 7.5 cm from the start position, the visual target was smoothly moved away from the centre location by 1 of 7 amplitudes (including 0) with a minimum jerk trajectory over the next 7.5 cm, before moving back to match the hand position with a minimum jerk trajectory over the next 7.5 cm of movement. Here participants need to ignore the visual discrepancy which is applied to the target. Again, all sensory discrepancy trials were performed in the null field such that participants could easily correct their hand motion as desired. In this protocol, participants should ideally correct for cursor perturbations but ignore target perturbations if they are able to independently control these feedback responses.

#### Session Organization

Each session in the two experiments was organized in a similar manner. Each session (which was performed twice for each experiment) consisted of 20 blocks of 70 trials, resulting in 1400 trials/session, and 2800 trials total for each experiment. Each block of seventy trials consisted of one of each of the 14 probe trials (all performed on channel trials) and 56 freely moving trials (in the null field) in which the cursor or target underwent a task-relevant or task-irrelevant motion as described above in the experimental protocols for each experiment. Specifically, there were four repetitions of each sensory discrepancy movement, which totalled 28 cursor movements where the target remained stationary, and 28 target movements where the cursor matched the hand movement. Within each block the trial order was fully randomized. Importantly the probe trials, which we use to assess any changes in the visuomotor feedback gains, were identical in both experiments and evenly applied throughout the entire experiments. A total of 40 repetitions for each probe trial type was applied in each experiment. Participants were required to take short breaks every 200 movements throughout the experiment. They were also allowed to rest at any point they wished by releasing a safety switch on the handle.

### Data Analysis

Analysis of the experimental data was performed with MATLAB R2024b. Position, velocity and endpoint force were low-pass filtered at 40 Hz with a tenth-order, zero phase-lag Butterworth filter. Lateral hand acceleration was calculated by differentiating the filtered velocity signal and then low-pass filtering at 40 Hz with a tenth-order, zero phase-lag Butterworth filter.

#### Adaptation to the Visual Task

To examine the adaptation to the two different conditions, we calculated a range of measurements across the experiment to compare the differences between the task-relevant and task-irrelevant sensory discrepancy environments.

##### Movement Duration

The movement duration for each trial was calculated as the duration between the cursor moving outside of the starting circle and entering the target circle. Mean and standard error of the mean across participants were calculated for each block of 20 trials.

##### Success Rate

The success rate was calculated as the percentage of trials in which good or great feedback was provided as defined above. Mean and standard error of the mean across participants were calculated for each block of 20 trials.

##### Peak Speed

The peak forward speed was calculated as the maximum velocity in the y-axis of the reaching movement between the start and end of the movement. Mean and standard error of the mean across participants were calculated for each block of 20 trials.

##### Max Lateral Motion

The maximum absolute lateral motion in response to the visual discrepancy was calculated on each non-probe trial. Trials in which the cursor underwent the sensory discrepancy were analysed separately from those in which the target underwent the sensory discrepancy. Mean and standard error of the mean across participants were calculated for each block of 20 trials.

##### Lateral Motion

To examine the timing, direction and magnitude of the hand motion to the imposed sensory discrepancies of the target and cursor during the movement, the lateral acceleration of the hand (x-axis) was examined for each type and magnitude of the sensory discrepancies. The lateral position was plotted as a function of the time. Mean and standard error of the mean were calculated for each time point in the movement. To examine the timing of the corrective responses, the mean hand acceleration was quantified over both an early (180-230 ms) and late (230-300 ms) time interval which corresponds to two previous windows of interest ^22,23,28^.

#### Rapid Visuomotor Responses

To measure the visuomotor responses to an imposed visual perturbation we used the lateral force measurements (x-axis) on the probe trials, in which participants moved within a mechanical channel. We first subtracted the mean x-axis force calculated between −200 to −50 ms prior to the start of each movement (when the hand was stationary) from every trial. This allows us to remove the effects of slight changes of the posture of the arm from influencing the forces measured in a trial. Individual probe trials were then aligned on visual perturbation onset. We then calculated the baseline response to the zero-visual perturbation using both the zero-cursor and zero-target probe trials (80 repetitions). This baseline response in each experiment was calculated as the mean lateral force (x-axis) at each time point across these 80 trials. For each of the fourteen different visual probe trials (7 sizes of cursor or target perturbations) we then calculated the force response to the visual probe as the response on that trial minus the baseline response on the zero-perturbation trials. This provides a direct measure of the motor response to the visual perturbation for each probe trial. To examine the feedback gain, we calculated the average post-perturbation force over two intervals: the first corresponding to a rapid involuntary response (180–230 ms) ^22^, and the second to a slower response (230-300 ms) ^23,28^.

To examine adaptation of the visuomotor feedback responses throughout the experiment we calculated the mean force response for each magnitude and type of probe trial within the two different intervals for each block throughout the experiment. The mean and standard error of the mean responses were then compared across the forty blocks to examine whether and how quickly the visuomotor feedback gains were adapted to the different sensory discrepancies that were applied differentially to the cursor and target during the movements. We then examined the change in the relative gain of the cursor and target feedback responses block by block through the experiment (for both intervals). The ratio was always calculated as the task-irrelevant response divided by the task-relevant response for appropriate contrast between the two experimental conditions. More specifically, we found the scaling factor that solved the linear equation between the magnitudes of the response to cursor and target perturbations for each size (mldivide). To examine the mean across participants, this was done for each participant using the data for all perturbation sizes for each block. To examine the scaling for each size perturbation, this was calculated across all the participants for each block.

To examine the visuomotor feedback responses after adaptation to the sensory discrepancies, the responses to visual perturbations in the first ten blocks of each experiment were not used for analysis to avoid the possible influence of initial high-gain or low gain trials at the beginning of the experiments. This choice was further supported by the results of the adaptation of the visuomotor feedback gains, which appeared to occur primarily within the first five blocks of each experimental protocol. Therefore, the visuomotor responses were measured as the responses on the last thirty blocks (*blocks 11–40*) of each experiment. To quantify the visuomotor gain, the mean responses were calculated for each probe trial magnitude in both the early and late intervals.

Finally, to examine whether there was a relative difference in the scaling of the visuomotor responses between the task-relevant and task-irrelevant sensory discrepancies, we calculated the ratio between the visuomotor responses in the task-relevant environment to the visuomotor responses in the task-irrelevant environment. As done previously, the ratio was always calculated as the task-irrelevant response divided by the task-relevant response for appropriate contrast between the two experimental conditions. The scaling factor was determined by solving the linear equation between the magnitudes of the response to cursor and target perturbations for each size (mldivide).

### Statistics

Statistics were performed in Matlab. Statistical significance was considered at the p<0.05 level for all statistical tests. Paired t-tests were performed to examine any differences between the initial 100 trials and final 100 trials on the success rate, movement duration and peak speed. Statistical differences across mean values and individual participants were done by calculating the 95% confidence intervals and examining whether the values overlap with each other or 1.0 which would indicate equal scaling between cursor and target responses. Where appropriate, t-tests were performed to determine whether the scaling values were equal to 1.0.

